# Assessment of dependency and consumption pattern of different forest products by the forest fringe villages of Shivalik Himalaya, Uttarakhand, India

**DOI:** 10.1101/2021.04.21.440776

**Authors:** J.S. Jiju, Soumya Dasgupta, Amit Kumar, Mohit Gera

## Abstract

Forest are essential for human beings for the enormous services it gives for livelihood and subsistence in the developing countries. We estimated the consumption and extraction levels of three major forest products viz., timber, fuelwood, and fodder in 20 forest fringe villages of the Timli forest range of Uttarakhand, India. We used a questionnaire-based household-level survey to collect information on the household economy and dependence of 380 households selected through stratified random sampling. We estimated that 69% of the overall yearly timber consumption of 20 villages, which comes around 2750 cubic meters (cum), was extracted from the nearby forest. The average timber consumption was 0.52 ± 0.22 cum household^-1^ year^-1^. We estimated the total annual fuelwood and fodder consumption to be 298913.89, and 204475 Quintal (Qt). The average fuelwood and fodder consumption were estimated to be 417.6 Quintal household^-1^ year^-1^ and 49 ± 9.1 Qt. household^-1^ year^-1^. We did general linear regression analysis to assess major biophysical and socio-economic determinants of villages and households for dependency on timber, fuelwood, and fodder. We found that the population of the village, distance from forest, distance from market, and annual average income are the major determining factors for timber, fodder, and fuelwood demand of the villages. Extraction of timber and non-timber forest products was the primary cause of depletion of forest biomass and forest carbon emission. Conservation effective management strategies in collaboration with all the stakeholder departments are needed to conserve forest resources with minimum extraction pressure from forest fringe villages of the study area.

**Highlights:**

- Questionnaire based household level survey was done to assess extraction and consumption patter of fuelwood, fodder and timber in Shivalik range of Uttarakhand, India.
- Fuelwood and Timber consumption was found high compared to other studies in the low altitude areas.
- 69% of total timber and 62% of the total fodder requirement met from the adjacent forest areas.
- Population of the village, distance from the forest and nearby market place, and annual average income were the major determinants of dependency of different forest products.

## 1. Introduction

Forest is an important natural resource for rural livelihood providing various services (Hussain et al 2019, Wunder et al 2014). Fuelwood is one of the oldest sources of energy known to man and used for over 500,000 years (Sharpe 1976), and by 2.7 billion people along with other traditional biomass, such as dung and agricultural by-products (IEA, 2010, 2017, Rahut et al 2016). In developing and underdeveloped countries, fuelwood is still the most important source of energy (Banyal et al 2013). Along with it, fodder and timber are the other forest products most commonly used by the rural people in developing nations (Singh and Sundriyal 2009, Nagothu 2001). More than one-third of the total energy demand, especially in the domestic sector of the developing countries’ rural population met through extracted biomass (Natarajan 1985; Vasudevan and Santosh 1987; FAO 2007). In developing countries, 70% of rural households use fuelwood as primary energy sources for cooking and space heating (Mishra 2008). Dependency on forest and other associated resources as the primary energy source were very high, especially in the rural areas of the developing countries (Hussain et al 2019). As for example, the dependency on forest biomass as primary source of energy was up to 87% in India (Madhu 2009, and Bhatt et al 2016), 77% in Nepal (Benato et al 2016), 78% in Bhutan (Rana et al 2016), 73% in Bangladesh (Huda et al 2014), 38.82% in Myanmar (Wen et al., 2017), 30% in Malawi (Fisher, 2004); up to 39% in western Ethiopia (Mamo et al., 2007); 40% in Zimbabwe (Cavendish, 2000) and up to 80% in Sub Saharan Africa (Sassen et al 2015). Fuelwood extraction often leads to forest degradation when the extraction is high, forest resources are limited, and alternative energy resources such as kerosene or Liquid Petroleum Gas (LPG) are unavailable (Kohl et al 2015, Specht et al 2015, WEC 2016, Nagothu 2001).

Agriculture and livestock rearing are significant livelihood sources in the villages situated near the forest areas (Kumar et al 2019). In turn, rearing livestock depends extensively on the forest, as forests are the major source of grass and fodder for livestock like bovine and ruminants reared by the villagers (Velho et al 2018, Nayak et al 2012). Open and uncontrolled grazing practices in Indian forests have an adverse impact on growing stock and regeneration (Agarwala et al 2016). Forest in India can support about 30 million livestock grazing, whereas 270 million cattle graze in (ICFRE 2001). According to Roy and Singh (2008), the estimated annual requirement and availability of dry fodder was to be 569 Metric Ton (MT) and 385 MT, and that of green fodder was to be 1025 MT and 356 MT, respectively. The difference in the requirement and availability clearly explains the high pressure on India’s forest due to the livestock population and how the demand for more fodder contributes to forest degradation in the country’s human-dominated landscapes.

In India, around 275 million people, coming around 40% of the country’s total poor population live in the forest fringes and depend on the forest for their livelihood (Velho et al 2018, Nayak et al 2012, World Bank 2006). On the other hand, more than 40% of the forests in the country are understocked and degraded (Aggarwal et al 2009; Bahuguna et al 2004). Gera et al (2017) revealed a loss of approximately 16% carbon from 1998 to 2014 due to forest degradation. Various factors starting from geographic to demographic and socio-economic are responsible for the degradation (Imai et al 2018, Rustagi et al 2010, Agarwal 2007). An increase in the agriculture depended population having low income, large and unproductive bovine population, restricted means of livelihood resulting in a vicious cycle of poverty exerting tremendous pressure on forests and make the ecosystem fragile to come back to its previous state (Davidar et al 2010). The adverse impacts of climate change will be additional pressure on the already vulnerable vegetation (Shrestha et al 2018, Chaturvedi et al 2011), and can significantly affect the flow of services from forests in terms of both quality and quantity (Nelson et al 2013; Scholes 2016). The impact will be on the forest dependent communities deriving their livelihood needs from the collection, handling, value addition, and selling of Non-Timber Forest Products, as the availability of many of the NTFPs is likely to be decreased more in the effect of climate change compared to timber and fuelwood (Robeldo and Forner 2005).

The socio-economic profile of an area determines the resource use pattern and enables the management authorities to prioritize the needs of the people inhabiting the area (Sharma et al 2009, Jain 2010, Barnes et al. 2011; Lee et al. 2013; Bansal et al. 2013). Additionally, in the current scenario, forest resources are insufficient, and forest degradation is happening at a faster rate (Gera et al. 2017). Effective human welfare and biodiversity conservation requires an understanding of household and village level factors determining the current patterns of forest resource dependencies and the potential changes in the future (Velho et al 2018, Ofoengbu et al 2017). It is crucial to evaluate the relationship between wealth and other socio-economic factors like livelihood patterns and socio-political assets modulating forest resources dependencies (Mascia et al 2014, Belchar et al 2015). In the present study, we estimated the extraction and consumption pattern of fuelwood, timber and fodder by the forest fringe villages of Timli reserve forest of Uttarakhand. We assessed the preferred species used as fuelwood and timber by the villagers. We also evaluated the demographic and socio-economic determinants of different forest resource extraction to understand the future conservation and developmental policy formulation for protection of diversity and integrity of the forest structure as well as decrease the resource dependency of the villagers of the study area.

## 2. Study area

The Shivalik landscape (29°57’ to 31°20’N and 77°35’ to 79°20’E) is the youngest of all the mountains in India and is significant biogeographically, as it has representative taxa from both Indo-Malayan to Palaearctic regions (Rawat and Mukherjee 2005). The fragile land formations, subtropical to a tropical climate, varied topography, and alluvial soil characterizes the region with undulating terrain intersected by seasonal streams (locally known as *Rau*) that drain this region (Johnsingh et al 2004; Rawat and Mukherjee 2005). We conducted the study in the forest fringe villages of the Timli Range of Shivalik region of Uttarakhand state, India (Figure 1). The study area is situated in the eastern part of Doon valley (30°19’ to 30°32’N and 77°34’ to 78°0’E) with an area of 99.07 km^2^, and falls under the Kalsi Soil Conservation Forest Division of Shiwalik Circle of Uttarakhand. The entire study area comprises 21 villages and 34 households (Gujjar Deras) of *Gujjar’s*, having a total population of 52,162 people (Census India data 2011). The Yamuna River bounds most of the forest area in the northwest and Shiwalik ridge in the south. The vegetation of this region mainly consists of tropical moist deciduous forests dominated by *Shorea robusta* and its major associates such as *Mallotus philippensis, Lagerstroemia parviflora, Ehretia laevis, Terminalia tomentosa* and plantations of *Tectona grandis, Eucalyptus* and Bamboo species. As per Champion and Seth (1968), Moist Shiwalik Sal forests (3C/C2a), Dry Shiwalik Sal Forests (5B/C1a), Northern Dry Mixed Deciduous Forests (5B/C2), and Low Alluvial Savannah Woodland (3/1S1) are the major forest types in the study area. The temperature of this region ranges from 40°C in summers to very cold (2°C) in winters (Yadav and Nandy 2015) and the average annual rainfall is 1550mm. Gujjar community, also known as Van Gujjars locally, are migratory pastoralists of the Himalayas and have distinct culture and traditions (Hussain et al 2017). Rearing buffaloes, goats and sheep are the primary source of income for Gujjars. Other major inhabitants of the study area include Garhwali and Nepali.

**Figure 1.**
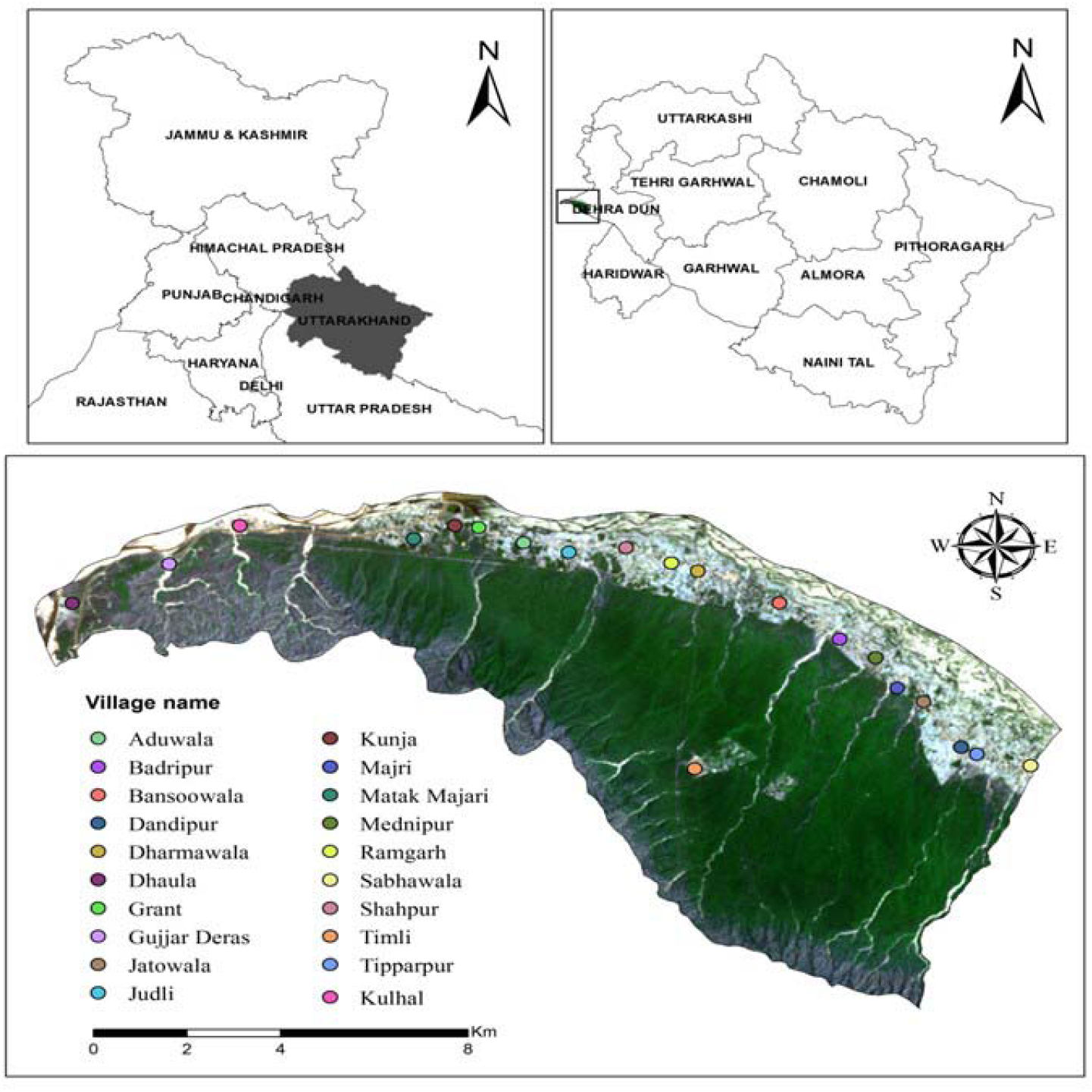
Map showing location of the villages in Timli Shiwalik region of Uttarakhand, India

## 3. Methodology

We carried out the study along with the working plan exercise of Indian Forest Service officers training courses of 2013-15 and 2014-16 during November 2014 and November 2015. Basic information collected from the Census India data set based on 2011 countrywide census data available in census.india website and verified through the village-level survey with the village head and elderly people of the villages. We have used s standardized questionnaire to conduct the socio-economic survey at the level of villages and households. The introduction on the village’s socio-economic profile i.e., population, occupation, landholding & land use, access to drinking & irrigation water, and other facilities such as school, hospital, government institutions, and market were recorded through focused group discussion at the village level. We recorded the information on the collection, use, and selling of NTFPs and the human-animal conflict during the village level discussion. After collecting socio-economic information at the village level, we collected information on household’s economy and dependence of households on forest products, viz., timber, fuelwood, fodder, grazing, and NTFP, etc through a household level survey. The population of the villages varied from 168 (*Gujjar Deras*) to 5200 (*Tipparpur*) individuals. We selected 15 to 25 households from each village for household-level surveys through stratified random sampling depending on the demography, economy, and societal status within each village. The information on household resource consumption and forest resource collection pattern was collected through interviews with a willing member of the household using a semi-structured questionnaire. We collected information on demography, occupation, fuelwood, timber and fodder dependency, and extraction pattern, livestock information, distance from the forest, and other resources like water for drinking and irrigation water source from each willing household member from 380 households. We estimated the mean values for the villages, from the information collected on fuelwood, timber and fodder collection and consumption. We also assessed the difference in fuelwood, timber, fodder consumption and the amount of extraction from the forest. The details of the household information and average consumption pattern of different forest resources given in Table 1.

**Table 1:** Number of villages and households surveyed in the socio-economic study with the Annual Timber Demand and Fodder Demand

| S.<br>No. | Name of<br>village | Number of<br>Households | Households<br>surveyed | Annual Timber Demand (CuM) |  |  |  | Fodder<br>quantity<br>(Qt) |
| --- | --- | --- | --- | --- | --- | --- | --- | --- |
|  |  |  |  | Housing | Furniture | Agri-Equipment's | Total |  |
| 1. | Grant | 450 | 25 | 25.56 | 15.84 | 1.98 | 43.38 | 26017.2 |
| 2. | Shahpur | 340 | 25 | 0 | 8.42 | 8.2 | 16.62 | 8190.6 |
| 3. | Judli | 428 | 25 | 32 | 6 | 2 | 40 | 4936.6 |
| 4. | Ramgarh | 250 | 15 | 1.87 | 6.55 | 0.67 | 9.101 | 14235 |
| 5. | Dharmawala | 400 | 25 | 26.55 | 2.38 | 1.75 | 30.68 | 15038 |
| 6. | Bansoowala | 55 | 15 | 5.92 | 9.76 | 0 | 15.68 | 10679.9 |
| 7. | Dandipur | 538 | 25 | 24.44 | 1.3 | 0.26 | 26 | 20107.9 |
| 8. | Kulhal | 110 | 15 | 45 | 30 | 5 | 80 | 4577.1 |
| 9. | Sabhawala | 400 | 25 | 17.39 | 2.2 | 0 | 19.591 | 34047.7 |
| 10. | Timli | 450 | 25 | 73.74 | 35.96 | 34.35 | 144.05 | 14125.5 |
| 11. | Kunja | 450 | 25 | 64.24 | 4.604 | 166.89 | 235.74 | 24273 |
| 12. | Aduwala | 190 | 25 | 48.28 | 40.8 | 88.4 | 177.48 | 9313 |
| 13. | Majri | 335 | 25 | 8.885 | 33.007 | 50.33 | 92.222 | 8005 |
| 14. | MatakMajri | 225 | 25 | 157.5 | 10.3 | 0 | 167.8 | 3580.6 |
| 15. | Mednipur | 750 | 25 | 28.8 | 8.34 | 1.05 | 38.19 | 3505 |
| 16. | Jatowala | 400 | 25 | 12 | 6.55 | 1.05 | 19.6 | 25700 |
| 17. | Dhaura | 65 | 15 | 12.88 | 1.786 | 6.908 | 21.574 | 652.9 |
| 18. | Tipparpur | 855 | 25 | 407 | 6 | 16 | 429 | 60115 |
| 19. | Gujjar<br>Deras | 168 | 15 | 777.6 | 318.4 | 0 | 1096 | 6329 |
| 20. | Badripur | 500 | 25 | 47.34 | 0.35 | 0 | 47.69 | 36354 |
| **Grand total** |  | **7359** | **450** |  |  |  |  |  |

We assessed the major fuelwood species used by the villagers and estimated the Use index value for all the species following Lance et al (1994) and Mitra et al (2017) using equation 1.

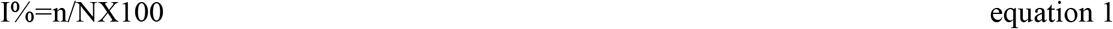

Use index (I%) presents the percentage of use, ‘n’ represents the number of villages citing the use of the tree species, and ‘N’ represents the total number of villages surveyed (Mitra et al 2017). The high value of the use index for a species denotes the high use of that particular species and vice versa.

We estimated the Pearson’s correlation coefficient of different factors affecting the annual timber, fuelwood, and fodder demand. We used generalized linear regression (GLM) analysis to identify the major determinants affecting the timber, fodder, and fuelwood extraction (response variables) by the household using R programming software (R version 3.4.4, 2018-03-15). Earlier studies revealed that education, income, household size, and access to clean energy like LPG availability determines the household’s choice of using forest biomass as energy source (Jain 2010, Lee et al 2013, Bansal et al 2013). Along with these factors, we used the village’s total population, distance from the forest, distance from the market, literacy, and net grazing requirement of the village as factors for the GLM analysis. We used the value of the corrected Akaike Information Criteria (AICc) to select the best model representing the major determinants of three response variables.

## 4. Results

We surveyed 20 forest fringe villages near the Timli forest division and the number of households varied from 34 in Gujjar Deras to 855 in Tipparpur village, and the mean household size estimated 7.1±0.17 persons. The overall literacy rate was 65.4%, which was lower compared to Dehradun city (84.2%), the state’s capital. *Dharmawala* (30%) and *Dhaula* (40%) had the least literacy rate. The primary source of livelihood includes wage labor and agriculture. However, individual families were also dependent on various occupations such as small-scale business, government, and private services for their livelihood. Wage labor is the primary source of livelihood for most of the respondents, followed by agriculture (18%), albeit small scale, business (7%), government or private services (4%), and others (4%); (Supplementary Figure 1). Within the *Gujjars*, the primary income source was pastoralism (90%), followed by wage labor (10%). The land holdings varied from landless to < 2 ha, however, most households (45%) have land holdings up to 0.5 ha, and 36.5% of the households, including *Gujjars*, are landless. The most common land use practice was agriculture (irrigated), wherein 64.5% of the land used to cultivate crops such as wheat, maize, millets, sugarcane, and rice. The other common land uses in the rest of the land was practiced for un-irrigated agriculture and horticulture. All the villages had education facilities within the vicinities of 1 km, except *Dhaula* (6 km), *Dandipur* (2 km), and *Tipparpur* (2 km) (Fig 1). Rearing livestock constitutes an essential source of livelihood. Most of the respondent families (90.4%) owned livestock; hence they were partially dependent on the adjoining forests for grazing and fodder collection. The total Adult Cattle Unit (ACU) holding in all villages was about 14240, with an average ACU of 2.03±0.26 per household. For grazing pressure on the forests, *Mednipur* village had the highest (2718) ACU and lowest (80.5) in *Aduwala* (supplementary Fig 2). The primary drinking water sources were tap water (46%) and hand pump (38%). The other sources like well (9.6%), spring or tank, or pond (6.4%) also contribute to drinking water accessibility. The major sources of irrigation were pump set (48.5%) followed by river or canal (36.5%), tank or pond (7%), and other water harvesting structures (8%).

### 4.1 Timber consumption pattern

We found that locals were dependent on the nearby forest to partially meet their timber demand for agricultural implements, cattle sheds, watch, and ward huts. The total consumption of timber was 991 cubic meters in the study area, of which 69% of the timber extracted from the nearby forest. Among the surveyed villages, Tipparpur, Timli, Majri, Dhaula, Gujjar Deras, Aduwala, and Jatowala were entirely dependent on the nearby forest for timber consumption (Table 1). The timber requirement was highest (66%) for housing, followed by furniture (26%) and agricultural implements (8%). The timber consumption and net timber requirement met from the forest were recorded highest (113 cubic meters; hereafter ‘cum’ and 74 cum, respectively) in *Badripur*, followed by *Sabhawala* (109 cum and 78 cum respectively), *Shahpur* (98 cum and 47 cum, respectively) and least in *Matak Majri* (9 cum and 6.5 cum, respectively) (Figure 2). The total annual timber consumption was 1929.9 cum. We recorded the average timber consumption to be 0.52 ± 0.22 cum household^-1^ year^-1^, and estimated the average per capita timber consumption to be 0.04 cum year^-1^ (Table 1).

**Figure 2.**
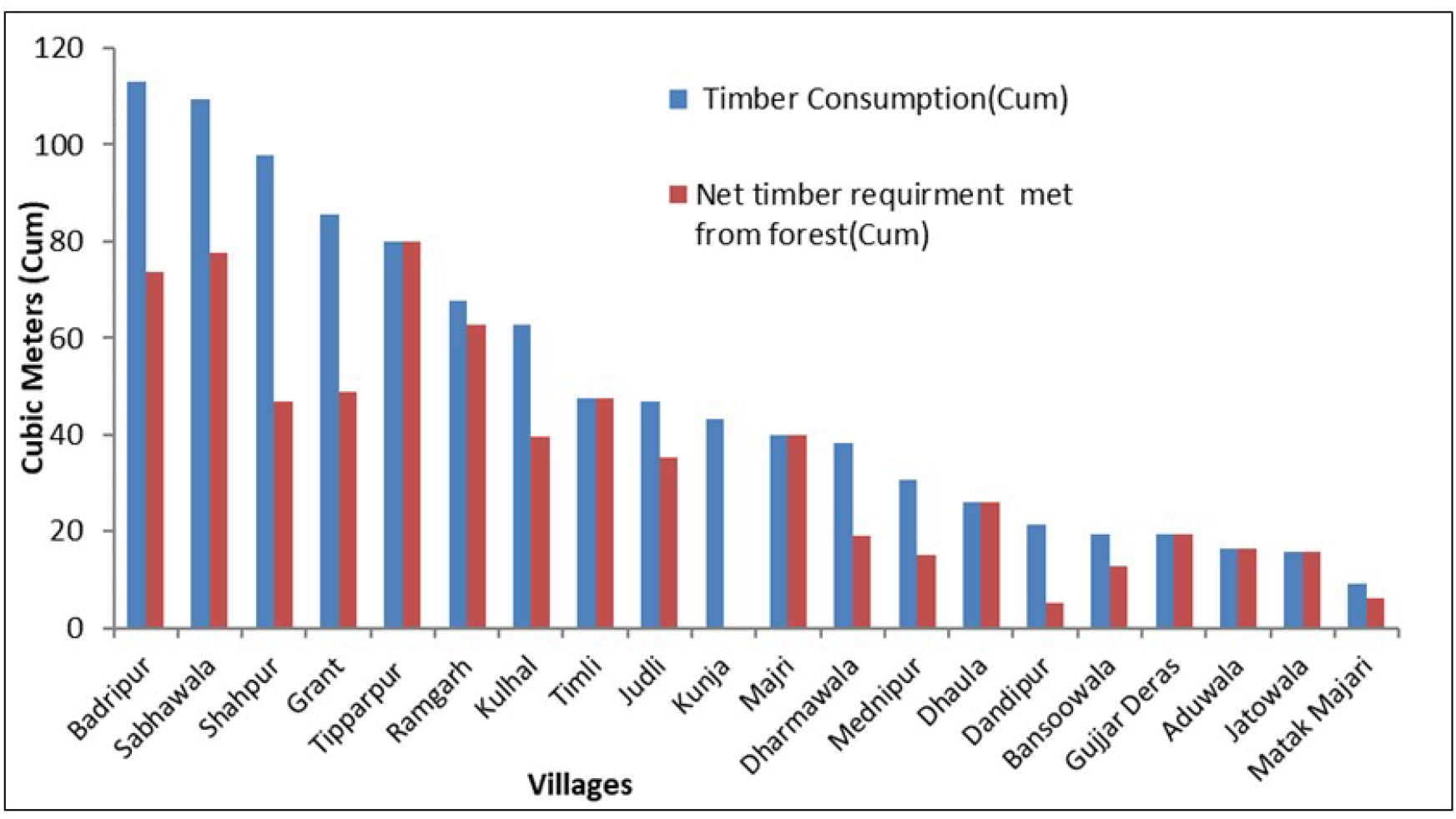
Timber demand and net Timber requirement met from forests in the Timli Shivalik region

### 4.2 Fuelwood consumption pattern

We found that villagers were mostly dependent on the forest for fuelwood, as 95% of the total fuelwood used to fulfill the household energy demand met from the fuelwood extracted from the nearby forests. About 89% of the fuelwood extracted was used for household energy demand, and 11% were sold to the nearby market for income generation. Fuelwood consumption and net fuelwood requirement met from nearby forests were highest in *Timli* (32499.6 Qt. and 30874.6 Qt., respectively) followed by *Mednipur* (30727.7 Qt. and 30113.2 Qt., respectively), *Dandipur* (29202.6 Qt. and 29202.6 Qt., respectively), and least in *Gujjar Deras* (1649 Qt. and 1649 Qt., respectively; Figure 3). The high fuelwood demand in Timli and Mednipur might be because of the low average income and high population of the villages. In *Gujjar Deras*, the fuelwood demand and net fuelwood requirement from nearby forests were least due to a low population (168 individuals). The total annual fuelwood consumption recorded was 29419.3 Qt. The average fuelwood consumption recorded was 41.1 ± 3.5 Qt. household^-1^ year^-1^ and in terms of average per capita fuelwood consumption was about 5.6 Qt. year^-1^. We found that, due to the inclusive dependency of *Gujjars* on the forests, the per capita fuelwood demand was high (9.8 Qt. year^-1^) as compared to total per capita fuelwood demand (5.6 Qt. year^-1^) (Figure 3).

**Figure 3.**
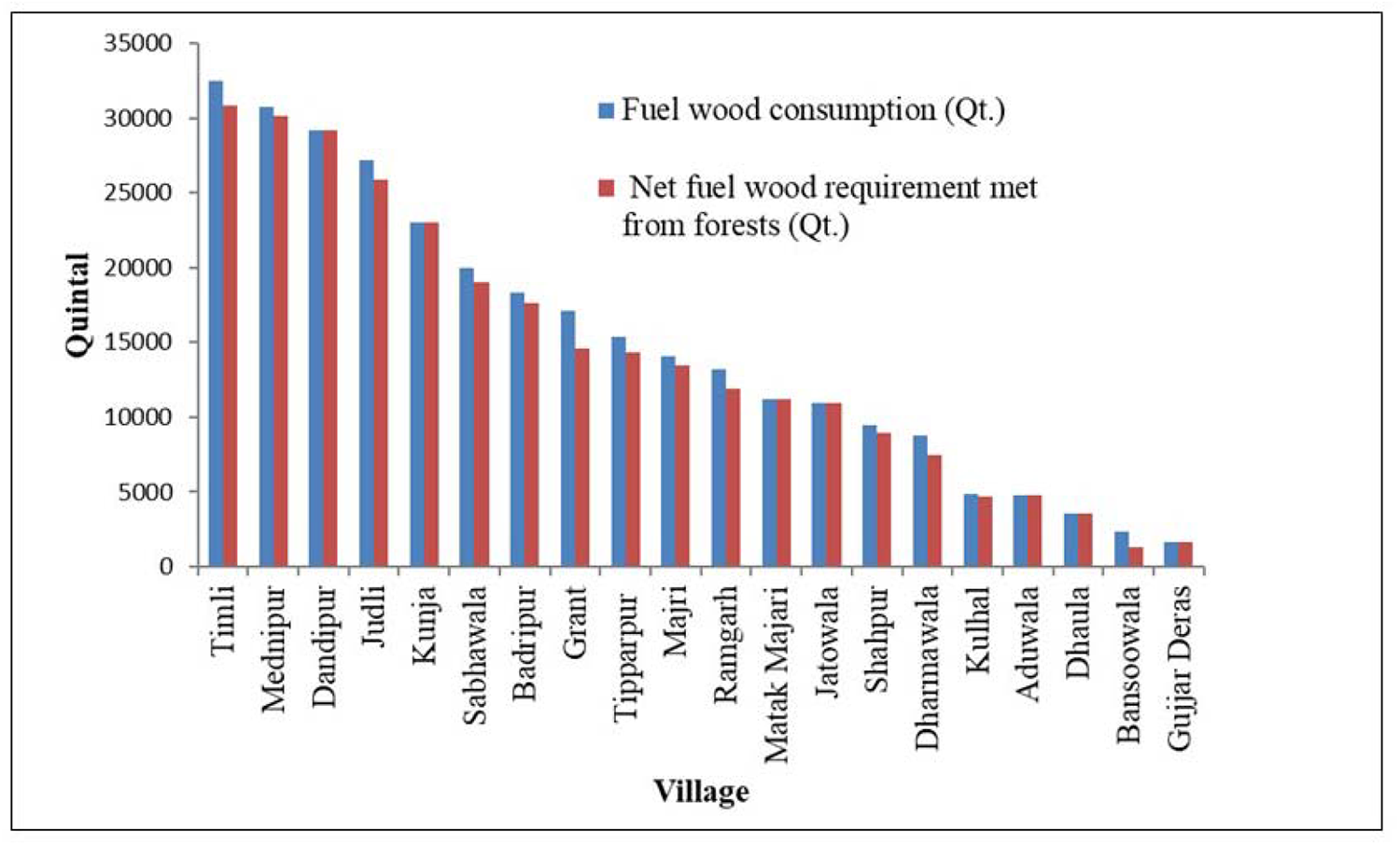
Fuelwood demand and Net fuelwood requirement (in Quintal) from Forest of studied villages.

### 4.3 Fodder consumption pattern

We recorded the total fodder extraction from the study area to be 204475 Qt with an average of 4949 ± 9.1 Qt. household^-1^ year^-1^. The average per capita fodder consumption estimated was 3.92 Qt. year^-1^. We found that the annual fodder consumption of Tipparpur village was the highest (60115 Qt.) followed by *Badripur* (36354 Qt.), *Sabhawala* (34047.7 Qt.), while *Dhaula* had the least (652.9 Qt.; Table 1). Per household fodder demand was highest in Bansoowala followed by Sabhawala and Badripur (Figure 4). The percentage of fodder demand met from nearby forests is highest in Gujjar deras and Timli followed by Dhaula and Kulhal (Figure 4). The locals obtain fodder primarily from nearby forests and farmlands, and from the roadside, riverside forest, and agro-forestry as per the availability. We recorded 17 species belonging to 14 genera and 10 families, primarily used as fodder by the local inhabitants. The major fodder species extracted from the forests consist of *Shorea robusta, Mallotus philippensis, Anogeissus latifolia, Millettia auriculata, Ehretia laevis, Grewia elastica, G. oppositifolia, Haldinia cordifolia, Desmodium oojeinense, Terminalia tomentosa, Bauhinia variegata, B. purpurea, Albizia lebbeck, Ficus benghalensis, F. racemosa, Lannea coromandelica*, and *Eulaliopsis binnata*. The important cultivated fodder crops grown in farms include *Trifolium alexandrinum* locally known as *barseem, Sorghum bicolor* locally known as *chari*, and residues of wheat, paddy, and maize.

**Figure 4.**
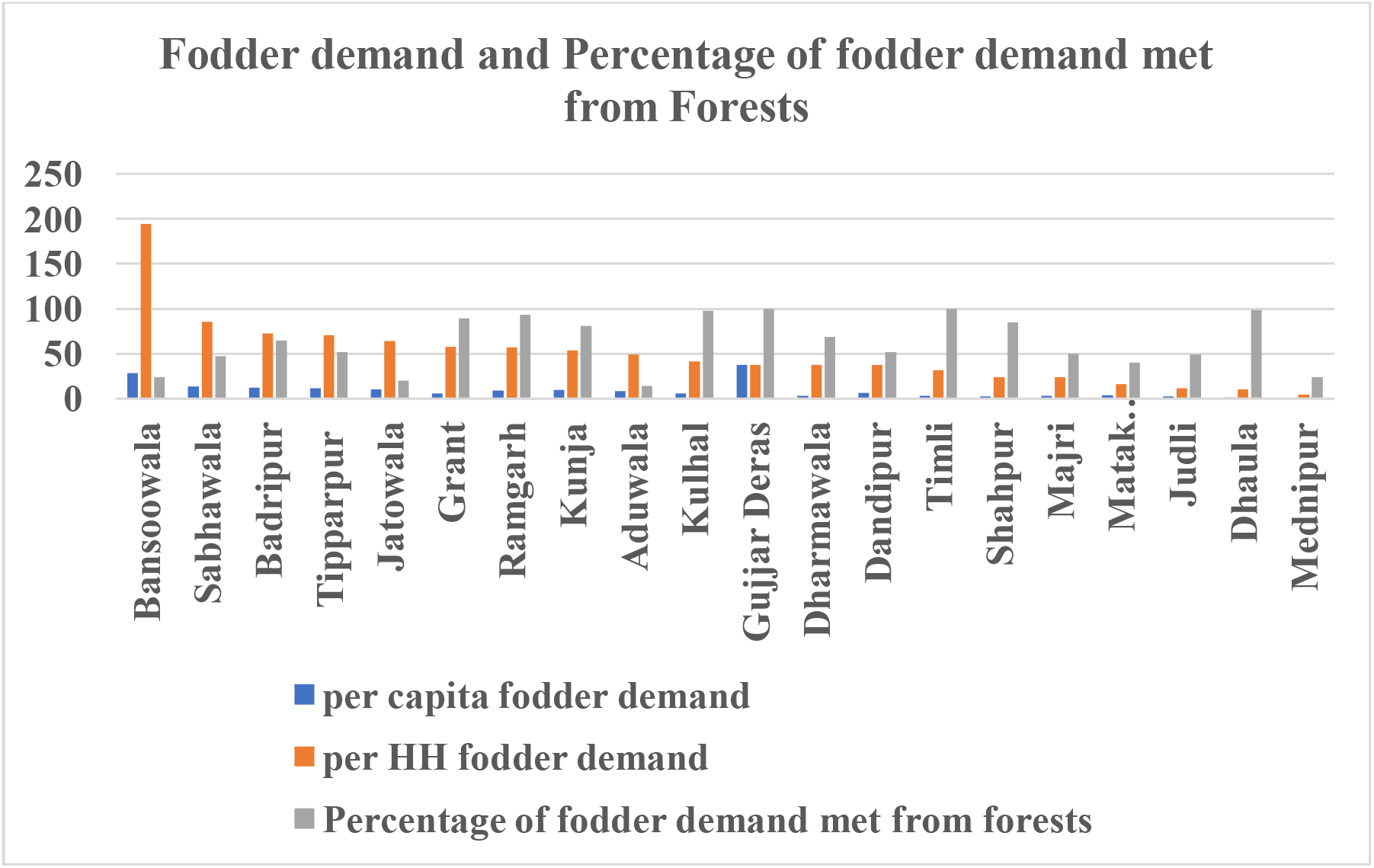
Per capita fodder demand, per household fodder demand (in Quintal) and percentage of fodder demand met from nearby Forest of studied villages.

### 4.4 Fuelwood use and preference

We found ten species belonging to ten genera and eight families were primarily used as fuelwood by the local inhabitants, mostly for cooking, boiling water, and space heating. We estimated the use-value of the species ranged from 5 to 95%, which was highest for *Shorea robusta* (95%; locally known as *Sal*) followed by the major associates of *Sal* forests *viz*., *Mallotus philippensis* (90%; locally known as *Rohini*) and *Terminalia tomentosa* (30%; locally known as *Sain*). The high use value indicates their great acceptability and availability as fuelwood, as well as, high anthropogenic pressure on these species. The remaining species include *Syzygium cumini, Anogeissus latifolia, Ardisia solanacea, Eucalyptus tereticornis, Tectona grandis, Clerodendrum viscosum*, and *Desmodium oojeinense* showed <15% use-value, reflected their low availability and low preference. The major fuelwood species with their ‘Use Value’ based on quality, characteristics, and availability in the area given in Table 2 and 3.

**Table 2.**
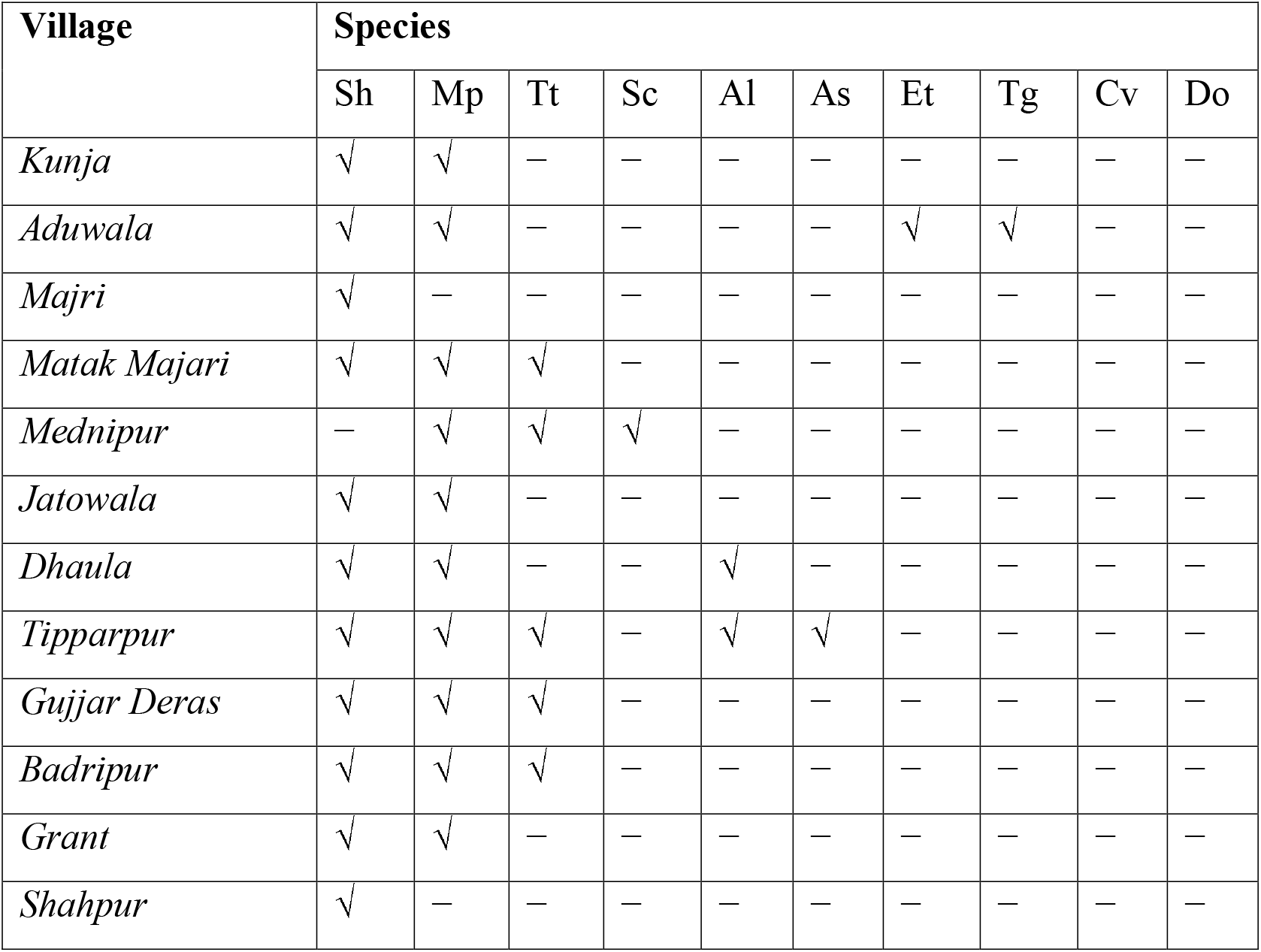
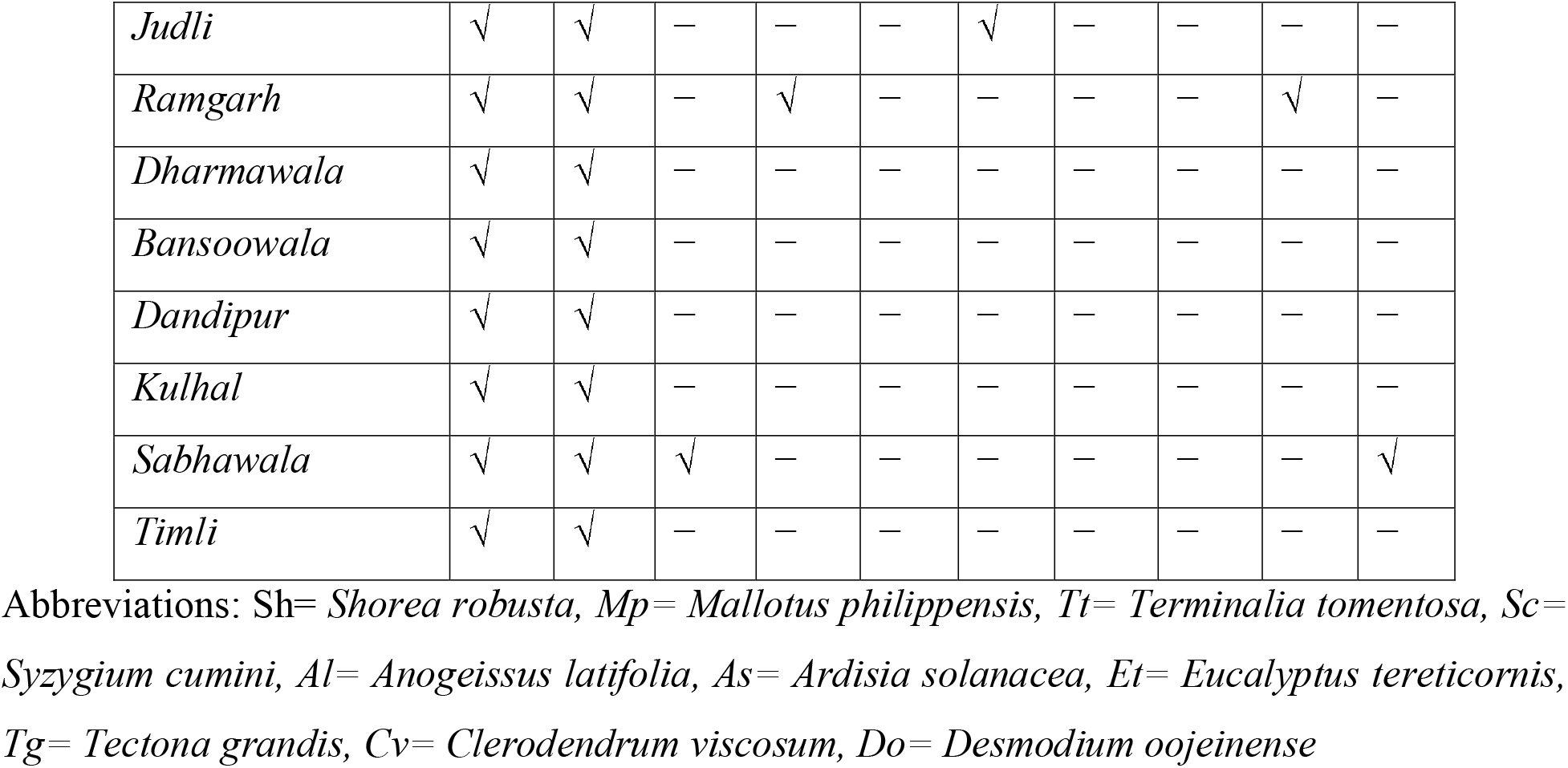
Major fuelwood species used in different villages of the Timli Range of Shivalik region, Uttarakhand, India.

**Table 3.** Major fuelwood species with their ‘Use Value’ based on quality, characteristics and availability in Timli Range of Shivalik region, Uttarakhand, India.

| Species | Family | Density (ind. ha<sup>-1</sup>) | Availability* | Use Value (%) |
| --- | --- | --- | --- | --- |
| *Shorea robusta* | Dipterocarpaceae | 146.1 | High | 95 |
| *Mallotus philippensis* | Euphorbiaceae | 34.4 | High | 90 |
| *Terminalia tomentosa* | Combretaceae | 5.2 | Medium | 30 |
| *Syzygium cumini* | Myrtaceae | 4.6 | Medium | 10 |
| *Anogeissus latifolia* | Combretaceae | 1.1 | Very low | 10 |
| *Ardisia solanacea* | Primulaceae | 0.8 | Very low | 10 |
| *Eucalyptus tereticornis* | Myrtaceae | 0.6 | Very low | 5 |
| *Tectona grandis* | Lamiaceae | 9.2 | Low | 5 |
| *Clerodendrum viscosum* | Verbenaceae | 0.5 | Very low | 5 |
| *Desmodium oojeinense* | Fabaceae | 1.8 | Very low | 5 |

\* Availability (individuals ha<sup>-1</sup>): Very low= <100, Low= 100-200, Medium= 200-300, High = 300-400, and Very high = >400. Species: a = tree, b = shrub.

### 4.5 Determinants of resource dependency and resource extraction

We found that annual fuelwood consumption, annual fodder consumption, and net requirement of grazing were positively correlated with the number of households and population of the villages (r^2^=0.714, 0.583 and 0.771, respectively). The percentage of fuelwood requirements met from the forest was negatively correlated with the household’s average annual income (r^2^=- 0.494). Distance from the nearby market and annual fuelwood demand was positively correlated (r^2^=0.557). We found that distance from the forest and average annual income were the significant determining factors that affects the household’s timber extraction pattern (Figure 5 and supplementary S2). We found the distance from the market, the average annual income of the villagers, and the population of the village were the significant determinants of fuelwood extraction by the households (Figure 6, summary given in supplementary S3). The GL models with fodder extraction pattern as response variable showed the population of the village, net grazing requirement, and fodder demand from the forest were the primary determining factor Figure 7 and supplementary S4). The details of all the GL models with parameters used and AIC values were given in Table 4.

**Table 4:**
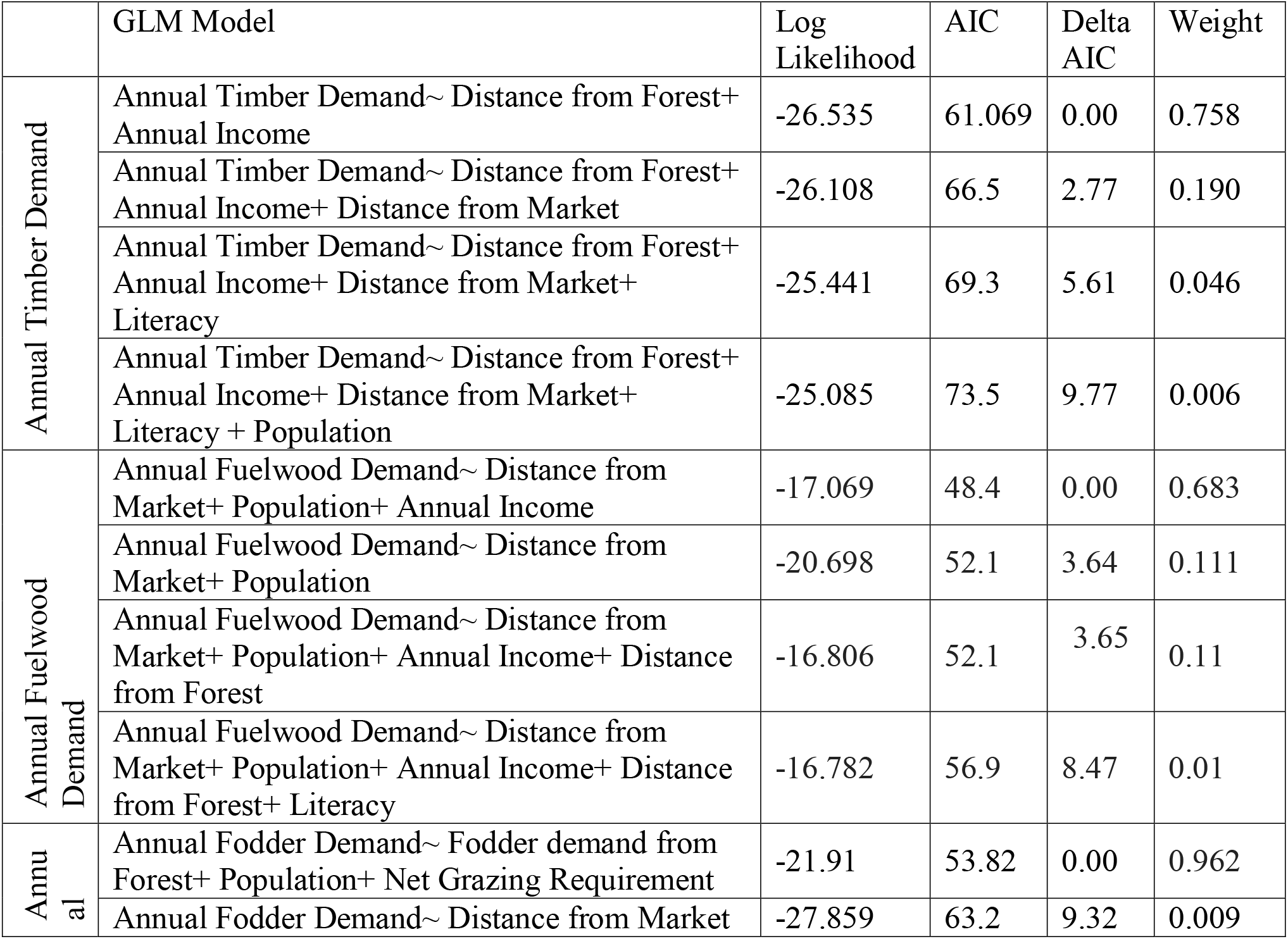
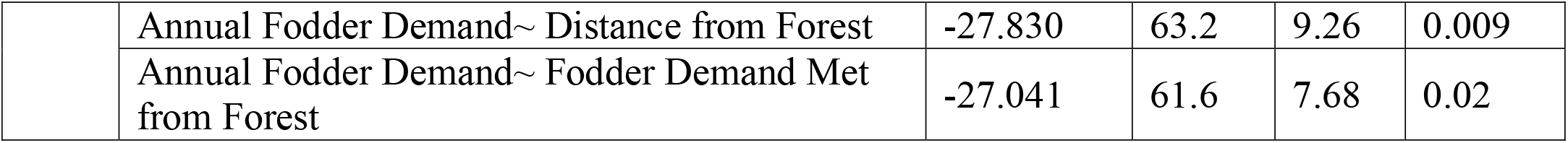
Table showing AIC values of different GLM

|  | GLM Model | Log Likelihood | AIC | Delta AIC | Weight |
| --- | --- | --- | --- | --- | --- |
| Annual Timber Demand | Annual Timber Demand~ Distance from Forest+ Annual Income | -26.535 | 61.069 | 0.00 | 0.758 |
|  | Annual Timber Demand~ Distance from Forest+ Annual Income+ Distance from Market | -26.108 | 66.5 | 2.77 | 0.190 |
|  | Annual Timber Demand~ Distance from Forest+ Annual Income+ Distance from Market+ Literacy | -25.441 | 69.3 | 5.61 | 0.046 |
|  | Annual Timber Demand~ Distance from Forest+ Annual Income+ Distance from Market+ Literacy + Population | -25.085 | 73.5 | 9.77 | 0.006 |
| Annual Fuelwood Demand | Annual Fuelwood Demand~ Distance from Market+ Population+ Annual Income | -17.069 | 48.4 | 0.00 | 0.683 |
|  | Annual Fuelwood Demand~ Distance from Market+ Population | -20.698 | 52.1 | 3.64 | 0.111 |
|  | Annual Fuelwood Demand~ Distance from Market+ Population+ Annual Income+ Distance from Forest | -16.806 | 52.1 | 3.65 | 0.11 |
|  | Annual Fuelwood Demand~ Distance from Market+ Population+ Annual Income+ Distance from Forest+ Literacy | -16.782 | 56.9 | 8.47 | 0.01 |
| Annual Fodder Demand | Annual Fodder Demand~ Fodder demand from Forest+ Population+ Net Grazing Requirement | -21.91 | 53.82 | 0.00 | 0.962 |
|  | Annual Fodder Demand~ Distance from Market | -27.859 | 63.2 | 9.32 | 0.009 |
|  | Annual Fodder Demand~ Distance from Forest | -27.830 | 63.2 | 9.26 | 0.009 |
|  | Annual Fodder Demand~ Fodder Demand Met from Forest | -27.041 | 61.6 | 7.68 | 0.02 |

**Figure 5.**
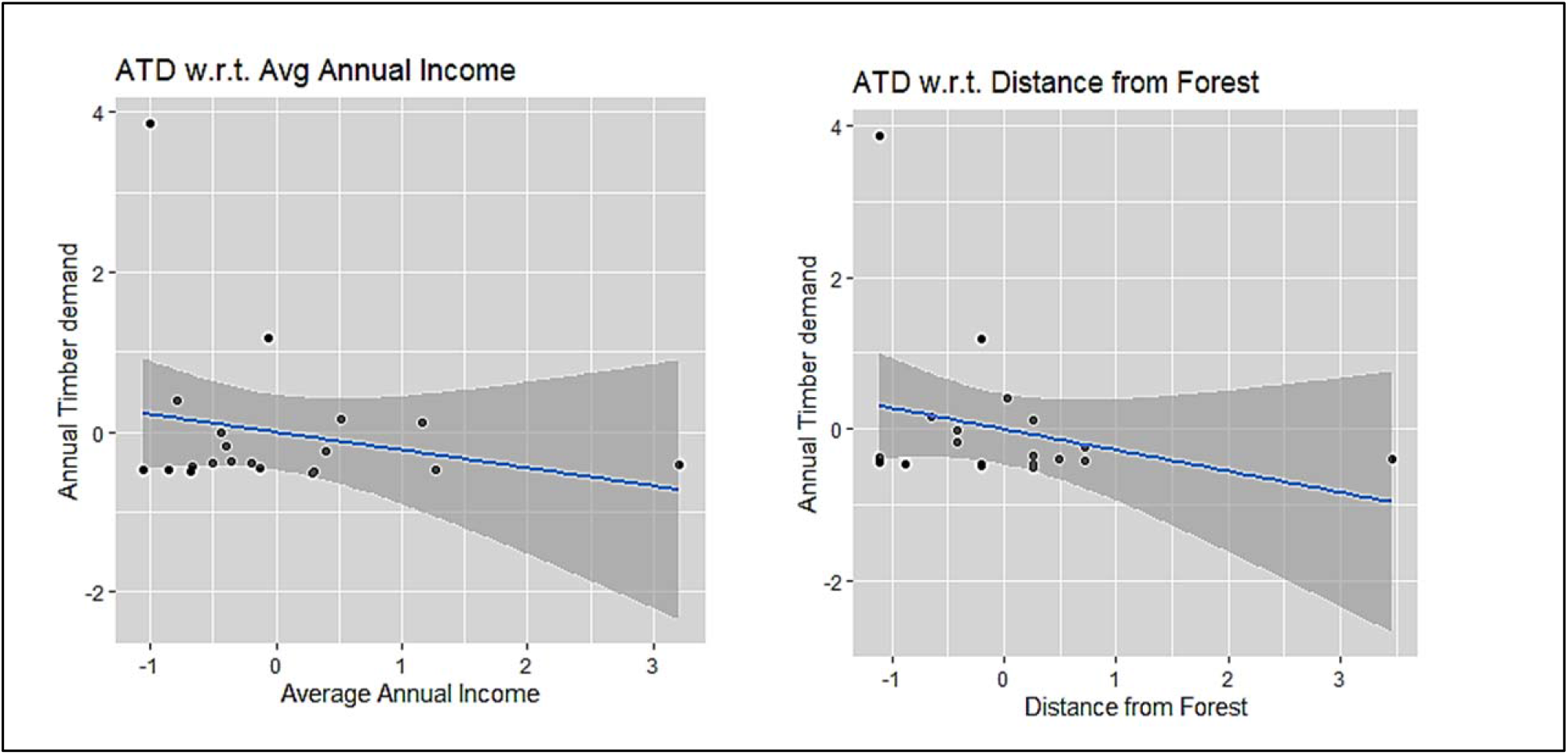
Annual timber demand with determining variables as found in generalized linear modeling

**Figure 6.**
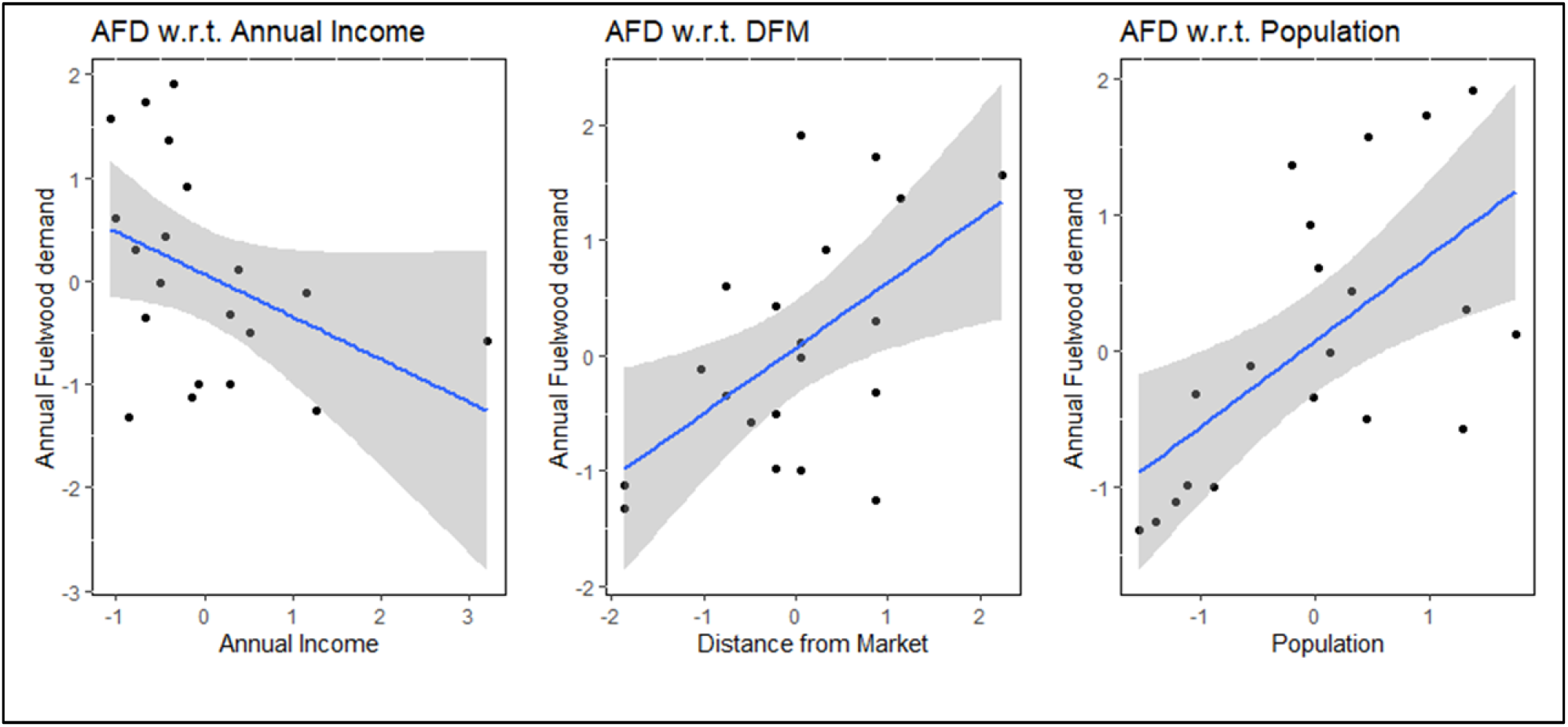
Annual fuelwood demand with determining variables as found in generalized linear modeling

**Figure 7.**
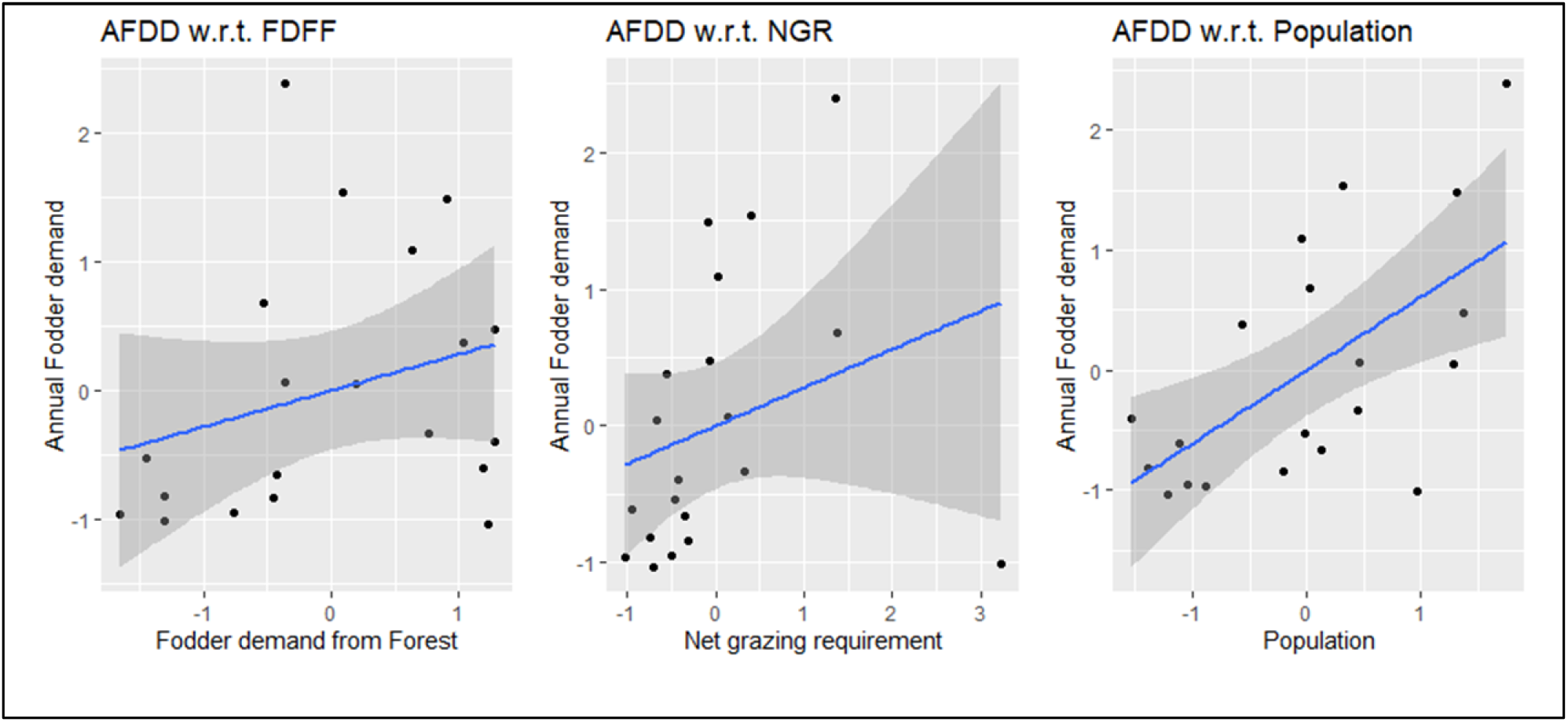
Annual fodder demand with determining variables as found in generalized linear modeling

## 5. Discussion

Energy is one of the primary requirements for social and economic development, and the demands vary regionally depending on the socio-economy and geographical conditions (Jain 2010, Lee et al 2013, Bansal et al 2013, Negi et al 2018). In developing countries like India, bio-fuel is a major energy source for people surviving at the subsistence level (Kumar and Sharma 2009; Negi and Maikhuri 2016). The fuelwood demand of the country ranges from 96–157 million tons having the consumption rate up to 148–242 kg per capita (Bhattacharya and Nanda 1992). However, annual consumption was estimated relatively high in various parts of the Himalaya (Campbell and Bhattarai 1984; Singh 1989; Metz 1990, Rawat et al. 2009; Negi and Maikhuri 2016; Bhatt et al. 2016). Poor accessibility of the alternative fuelwood sources makes the rural population entirely dependent on wood sources (Bhatt et al 2004). In most cases, the fuelwood demand is met solely from the adjoining forests (Hussain et al 2017). This uninterrupted extraction of fuelwood and fodder is the major reason for the depletion of forest patches (Singh 1998). Singh et al (2010) showed fuelwood consumption ranging from 20-25 kg households^-1^ day^-1^ in high-altitude areas of Garhwal Himalaya, Uttarakhand. Awasthi et al (2003) reported 14.65 kg households^-1^ day^-1^ fuelwood consumption in other villages of the Garhwal Himalaya. However, the per capita use values were higher than the ones reported from villages of lower altitudinal ranges of Western Himalaya i.e. 1.49 kg capita^-1^ day^-1^ (Bhatt et al 1994), for southern India (1.9–2.2 kg capita^-1^ day^-1^; Reddy 1981) and the Himalayan range of Nepal (1.23 kg capita^-1^ day^-1^; Mahat et al 1987). The change in fuelwood consumption was also evident in different altitudinal gradients, as in higher altitude due to cold temperature, fuelwood consumption was 2-3-fold high compared to low altitude, due to essential warming needs (Bhatt and Sachan 2004). In our study we found that the fuelwood consumption by the villagers ranged from 2.6–19.7 kg household^-1^ day^-1^ with an average of 11.26 kg household^-1^ day^-1^. The amount was higher than the consumption pattern in low altitude areas (Bhatt et al 1994, Mahat et al 1987), and India’s average fuelwood consumption, i.e. 4.06 kg household^-1^ day^-1^ in rural areas as per the Centre for Development Finance (www.householdenergy.in), but comparable with the consumption pattern of high altitude areas of Western Himalaya (Awasthi et al 2003, Singh et al 2010 and Negi et al 2018).

We found that the average fodder extraction from forest by the household was 7.51 kg household^-1^ day^-1^, which was much less than that of recorded earlier from other areas of Uttarakhand. Dhyani et al 2011 recorded the fodder extraction by the households near Kedarnath Wildlife Sanctuary range from 62.4 to 80.4 kg household^-1^ day^-1^ and Dhanai et al 2014 recorded fodder extraction range from 56.64 to 72.48 kg household^-1^ day^-1^ near the Takoligad watershed. The reason may be because of less livestock unit per household in the present study area and the villagers also collect fodder from the agricultural field and roadside area and from the bank of small rivulets also. Therefore, the pressure exerted to the forest area for fodder was found much less than expected.

Forest, pastures, arable land, cattle, and human population are the five essential components in the hill ecosystem which are linked with each other in a series of dynamic relationships starting from the production to transfer and consumption of the energy (Sharma et al 2009, Khuman et al 2011). The availability of fodder, fuelwood, and litter is vital for the survival and livelihood of the rural settlements in the Himalayas (Dhyani et al 2011, Dhanai et al 2014). In most Indian Himalayan regions, fuelwood obtained from the forest is the sole source of energy available to the residents (Kumar and Sharma 2009; Negi and Maikhuri 2016; Thapa and Weber 1990). The present study reveals the population of the village, distance from the forest, distance from the market, and annual average income are the limiting factors for timber, fodder, and fuelwood demand of the villages. The studies from Barnes et al. 2011; Lee et al. 2013; Bansal et al. 2013, Jain (2010) and Bansal et al. (2013) from India; Arthur et al. (2010) from Mozambique; Andadari et al. (2014) from Indonesia Pine et al. (2011) from rural Mexico, Jan (2012) from northwest Pakistan, Beyene and Koch (2013) from urban Ethiopia concluded that the household size, education and household income are the most significant factors that determine the willingness to use cleaner energy instead of forest biomass. Subject to availability, fuelwood as an essential energy source is easy to collect and use (Specht et al 2015), whereas the other commercial sources of energy are beyond the reach due to accessibility in remote areas. Due to remoteness, alternative sources come with high prices and limited supply (FAO 2007). It was evident in various studies, that the rural population makes greater use of wood for heat and cooking fuel (Miah et al 2003; Moran-Taylor and Taylor 2010). So, in the past few years, there is an enormous amount of attention given for reducing biofuel use, as it is nested within the three major challenges of the developing world – energy, poverty, and climate change (FAO 2007). According to Kanagawa and Nakata (2007), fuelwood consumption increases the direct payments of rural households, and fuelwood collection also takes valuable time and effort resulting in loss of education and income generation opportunity for collectors (Hussain et al 2017). Unsustainable fuelwood collection and inefficient conversion technology used in remote rural areas have severe implications on the environment (Arnold et al 2003; Chen et al 2006). In recent years, scientists and planners across the world have shown concern about the gap between forecasted estimates of biomass supply and demand. The increased demand will place stress on women, children, and the environment, at the national level and different eco-zones of the country (Gadgil et al 1989).

The estimation of extraction and consumption pattern of major forest products in the remote rural area is crucial as the human population is increasing and the socio-economic pattern and the environmental conditions are changing day by day. The mean household size of the studied village is higher than the mean household size of the communities living in the hilly areas of Uttarakhand and that of the whole country average of 5.3 persons (Census of India 2011). The majority of respondents (67%) indicated wage labor as the primary source of livelihood, which directly indicates that the economic status of the major population is low, which has resulted in a low literacy rate (65.4%) as compared to Dehradun (84.2%). The low literacy rate and employment of the locals can be improved with the support for better educational and employment opportunities.

As the vegetation of the study area is dominated by *Shorea robusta* and its major associate timber species such as *Mallotus philippensis, Anogeissus latifolia, Syzygium cumini*, and *Terminalia tomentosa*, the pressure in terms of the utilization of these resources by the locals is higher as compared to the other species. Practice of selective species harvesting has already reported to affect the species composition and assemblage (Negi and Maikhuri 2016, Rawal et al 2012, Singh et al 2010). Therefore, to reduce the anthropogenic pressure on the forests, the plantation of indigenous and multipurpose species in consultation with the local inhabitants and the concerned forest department would be the most practical solution. To reduce forest dependency and reduction in forest degradation, policy measures to be taken by different stakeholders. The use of LPGs and their availability to the villagers to be taken care of through regular supply at a subsidized price. The other suggestive mitigation measures could be the creation of Self Help Groups for alternative livelihood options, assistance in cattle breed improvement and encouraging stall feeding, the establishment of small scale industries such as mushroom cultivation, leaf plates using *Sal* leaves, manure making, honey production, etc., along with initiation and encouragement of the locals for adopting biogas plants and solar power are few measures that need to be exploited for the fulfillment of the energy demand of the region. The National Mission for Enhanced Energy Efficiency (NMEEE) under India’s National Action Plan for Climate Change (NAPCC) is already working on these issues, but the inter-sectoral linkages and deficiency of data sets on energy requirement and consumption pattern of remote areas causing the difficulty implementing policy intervention in the remote areas.

## 6. Conclusion

Understanding the drivers for forest degradation is essential for developing policies and measures that aim to change the current trends in consumption patterns of forest products towards a more environmentally friendly outcome. In the present study, we found that lack of adequate education and dependency of the residents on the natural resource-based livelihood resulting in the high dependence on the timber and non-timber forest resources and thus high level of extraction of forest products from the Timli forest. We found that the fuelwood extraction pattern was high when compared to other published literature in low altitude areas, whereas the fodder collection is much less compared to other study result in Uttarakhand. The high level of fuelwood and timber extraction will degrade the forest resources in the future and is not a good option for the future sustainability of the area and its residents. Policy level intervention is needed urgently for alternative livelihood opportunities for the people and to ensure alternative energy sources for cooking and other daily needs. The plantation can be a sustainable option for the rejuvenation of forest unless and until there is a change in resource extraction for household and economic needs. Integration of different stakeholders’ activity for reducing forest dependency and generating alternative sustainable livelihood for the improvement of the socio-economy of the area is urgently needed to conserve the last remaining forest patch and the diversity of the study area for its sustainable future.

## Acknowledgements

The authors gratefully acknowledge Sh. Vinod Kumar, former Director and Dr. Shashi Kumar, Director of Indira Gandhi National Forest Academy (IGNFA) for consistent support and guidance. We also wish to acknowledge Dr. Jagdish Kishwan, Chairman *‘Apex Academic Committee on REDD-plus in relation to global warming and Climate Change’* for kind support during various meetings of REDD-plus Cell at IGNFA. The authors are also indebted to IFS Probationers of 2014-16 and 2015-17 courses for their participation in the smooth conduct of the socio-economic survey. Thanks, are also due to Dr. Taibanganba Watham for his help in preparation of the map.

## Author Contributions

Conceptualization and study design (Jiju J. S., Dr. Mohit Gera); Methodology Formal Analysis and Investigation (Jiju J. S., Soumya Dasgupta, Amit Kumar); Original Draft Preparation (Jiju J. S. and Soumya Dasgupta); Review and editing (Soumya Dasgupta, Amit Kumar and Mohit Gera); Resources (Dr Mohit Gera and Jiju j. S.) Supervission (Dr Mohit Gera)

## Permission and informed consent

All the necessary permission and informed consent was taken by the authors wherever it is needed, for the study.

