## Supplementary File for "Assessment of dependency and consumption pattern of different forest products by the forest fringe villages of Shivalik Himalaya, Uttarakhand, India"

Supplementary table 1 - DATA DETAIL TABLE

|  | **Household-level** | Data source | Data used |
| --- | --- | --- | --- |
| 1 | Education | Household-level survey | Ranked |
| 2 | Occupation | Household-level survey | Ranked |
| 3 | Total members earn | Household-level survey | Number |
| 4 | Avg. Household size | Household-level survey | Number |
| 5 | Species Preference for fuelwood | Household-level survey | Ranked |
| 6 | Annual Timber Demand (CuM) | Household-level survey | Number |
| 7 | Annual Fodder Demand (Qt) | Household-level survey | Number |
| 8 | Annual Fuelwood demand (Tonnes) | Household-level survey | Number |
| 9 | Grazing in (ACU) | Household-level survey | Number |
| 10 | LPG connection | Household-level survey | Presence/absence |
| 11 | Total family member | Household-level survey | number |
| 12 | Avg Land Holding per HH (ha) | Household-level survey | Number |
| 13 | Human-Animal Conflict | Household-level survey | Ranked |
| 14 | Source of Drinking water | Household-level survey | Ranked |
| 15 | Source for irrigation | Household-level survey | Ranked |
| 16 | Distance to Nearest Market for forest Products | Household-level survey | Ranked |

S 2. Summary of the GLM model Annual Timber Demand~Distance from Forest +Annual Income.

Deviance Residuals:

| Min | 1Q | Median | 3Q | Max |
| --- | --- | --- | --- | --- |
| -0.7798 | -0.5023 | -0.3709 | 0.2860 | 3.3443 |
| Coefficients | Estimate | Std. Error | t- Value | Pr(>t) |
| Intercepts | 9.727e-11 | 2.212e-01 | 0.000 | 1.000 |
| DFF | -2.731e-01 | 2.270e-01 | -1.203 | 0.245 |
| AI | -2.185e-01 | 2.270e-01 | 0.963 | 0.349 |
| (Dispersion parameters for gaussian family taken to be 0.) | | | | |
| Null deviance: 19.000 on 19 degrees of freedom | | | | |
| Residual deviance: 16.632 on 17 degrees of freedom | | | | |
| AIC: 61.069 | | | | |

S 3. Summary of the GLM model Annual Fuelwood Demand~Distance from Forest+Population+ Annual Income

| Min | 1Q | Median | 3Q | Max |
| --- | --- | --- | --- | --- |
| -1.08906 | -0.40166 | -0.01614 | 0.44368 | 0.94961 |
| Coefficients | Estimate | Std. Error | t- Value | Pr(>t) |
| Intercepts | 0.06809 | 0.14202 | 0.479 | 0.63812 |
| DFM | 0.34997 | 0.15505 | 2.257 | 0.03833* |
| Population | 0.53993 | 0.15389 | 3.508 | 0.00291** |
| AI | -0.39035 | 0.14754 | -2.646 | 0.01762* |
| Significance codes: 0’***’, 0.001’**’, 0.01’*’, 0.05’.’, 0.1’ ‘ | | | | |
| (Dispersion parameters for gaussian family taken to be 0.) | | | | |
| Null deviance: 19.7618 on 19 degrees of freedom | | | | |
| Residual deviance: 6.4546 on 16 degrees of freedom | | | | |
| AIC: 44.139 | | | | |

S 4. Summary of the GLM model Annual Fodder Demand~Percentage of fodder demand met from forest + Population + Net requirement of grazing for the village

| Min | 1Q | Median | 3Q | Max |
| --- | --- | --- | --- | --- |
| -1.3236 | -0.6102 | -0.1008 | 0.4421 | 1.4208 |
| Coefficients | Estimate | Std. Error | t- Value | Pr(>t) |
| Intercepts | 1.095e-10 | 1.809e-01 | 0.000 | 1 |
| FDMF | 2.829e-01 | 1.944e-01 | 1.456 | 0.1648 |
| Population | 5.882e-01 | 2.228e-01 | 2.640 | 0.0178* |
| NGR | 3.555e-02 | 2.297e-01 | 0.155 | 0.8790 |
| Significance codes: 0’***’, 0.001’**’, 0.01’*’, 0.05’.’, 0.1’ ‘ | | | | |
| (Dispersion parameters for gaussian family taken to be 0.) | | | | |
| Null deviance: 19.000 on 19 degrees of freedom | | | | |
| Residual deviance: 10.474 on 16 degrees of freedom | | | | |
| AIC: 53.82 | | | | |

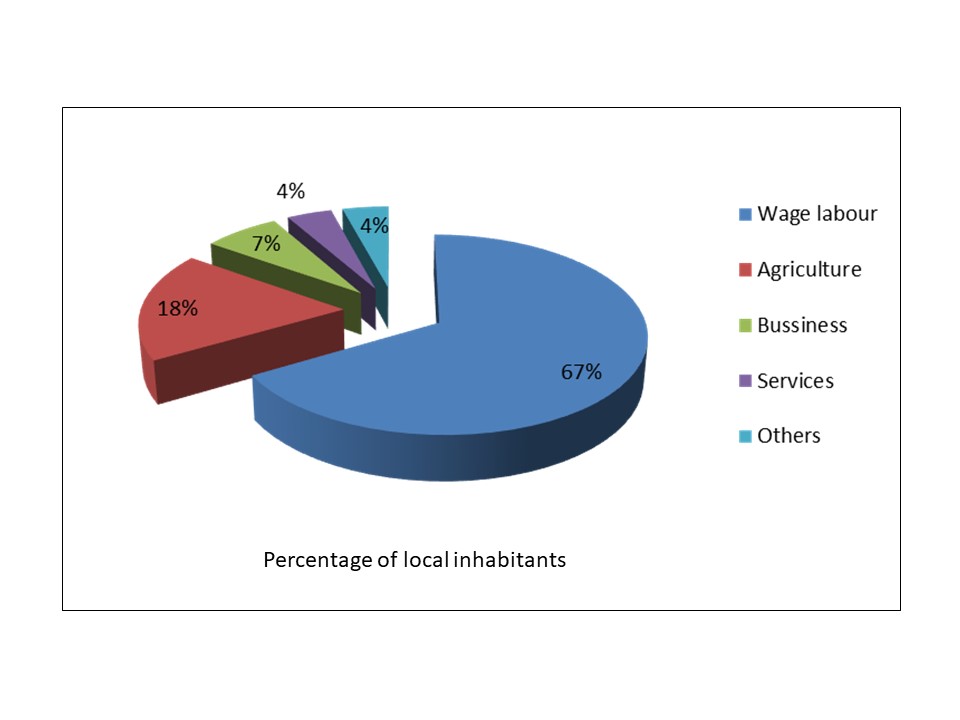

Supplementary Figure 1. Occupation of the local inhabitants in the Timli Range of Shivalik region.

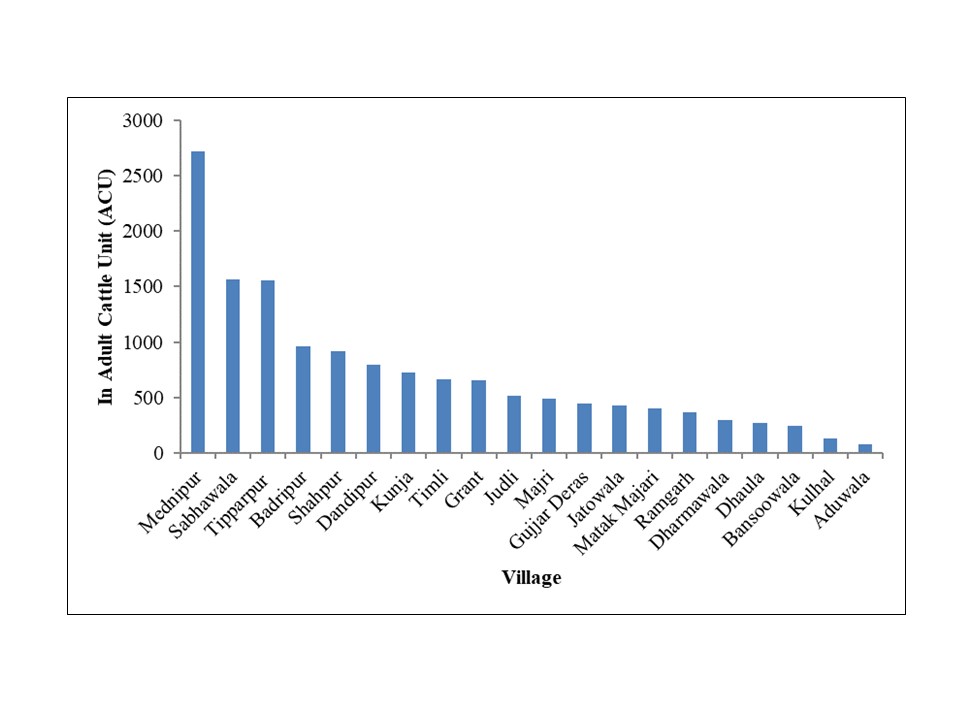

Supplementary Figure 2 – Total adult cattle unit (ACU) in the villages surveyed.
